# Changes in the Type 2 diabetes gut mycobiome associate with metformin treatment across populations

**DOI:** 10.1101/2023.05.25.542255

**Authors:** Emily Van Syoc, Michelle Pistner Nixon, Justin D. Silverman, Yuhong Luo, Frank J. Gonzalez, Ilze Elbere, Janis Klovins, Andrew D. Patterson, Connie J. Rogers, Erika Ganda

**Affiliations:** Department of Biology, The Pennsylvania State University, University Park, PA 16802, USA; Department of Animal Science, The Pennsylvania State University, University Park, PA 16802, USA; One Health Microbiome Center, The Pennsylvania State University, University Park, PA 16802, USA; College of Information Sciences and Technology, The Pennsylvania State University, University Park, PA 16802, USA; Departments of Statistics and Medicine, The Pennsylvania State University, University Park, PA 16802, USA; Center for Cancer Research, National Cancer Institute, National Institutes of Health, Bethesda, MD 20892, USA; Latvian Biomedical Research and Study Center, Riga, Latvia; Department of Veterinary and Biomedical Sciences, The Pennsylvania State University, University Park, PA 16802, USA; Department of Nutritional Sciences, University of Georgia, Athens, GA 30602, USA

**Keywords:** Gut mycobiome, type 2 diabetes mellitus, gut microbiome, metagenomics

## Abstract

The human gut teems with a diverse ecosystem of microbes, yet non-bacterial portions of that community are overlooked in studies of metabolic diseases firmly linked to gut bacteria. Type 2 diabetes mellitus (T2D) associates with compositional shifts in the gut bacterial microbiome and fungal mycobiome, but whether T2D and/or pharmaceutical treatments underpin the community change is unresolved. To differentiate these effects, we curated a gut mycobiome cohort to-date spanning 1,000 human samples across 5 countries and a murine experimental model. We use Bayesian multinomial logistic normal models to show that metformin and T2D both associate with shifts in the relative abundance of distinct gut fungi. T2D associates with shifts in the Saccharomycetes and Sordariomycetes fungal classes, while the genera *Fusarium* and *Tetrapisipora* most consistently associate with metformin treatment. We confirmed the impact of metformin on individual gut fungi by administering metformin to healthy mice. Thus, metformin and T2D account for subtle, but significant and distinct variation in the gut mycobiome across human populations. This work highlights for the first time that oral pharmaceuticals can confound associations of gut fungi with T2D and warrants the need to consider pharmaceutical interventions in investigations of linkages between metabolic diseases and gut microbial inhabitants.

## INTRODUCTION

The gut microbiome is a complex ecosystem of billions of micro-organisms, but research on the associations between human diseases and the gut microbiome has traditionally focused on bacteria. Interest in non-bacterial, intestinal communities such as the fungal mycobiome has increased in recent years, spurring studies to describe the gut mycobiome across disease phenotypes and healthy individuals^1–3^. Intestinal fungi comprise a small proportion of gut microbial biomass^4^ and can be difficult to analyze with traditional next-generation sequencing approaches. Nevertheless, multiple reports in the last few years suggest that the gut mycobiome may contribute to gut homeostasis and can instigate inflammatory responses when dysregulated ^5–8^. Differences in fungal community diversity metrics in case-control study designs suggest a state of fungal dysbiosis^3,9^ analogous to bacterial dysbiosis that contributes to the development of metabolic diseases^10,11^. Indeed, a recent review described perturbed gut fungi in multiple disorders including irritable bowel syndrome, colorectal cancer, liver diseases, COVID-19 infection, obesity, Type 2 diabetes, and neurological disorders^3^. However, human cohort studies of the gut mycobiome have small sample sizes and individual reports yield conflicting evidence, particularly in Type 2 diabetes (T2D).

In 2022, 28.7 million Americans had diagnosed Type 1 or Type 2 diabetes^12^. In 2017, the direct healthcare costs of diagnosed diabetes in the United States was USD 237 billion, and are projected to reach USD 825 billion by 2030^13,14^. Perturbations in the gut bacterial microbiome are reported in human and animal models of T2D, which may contribute to sub-clinical inflammation and insulin resistance^10,15,16^. Metformin is the most widely prescribed antidiabetic drug^17^ and is known to modulate gut bacteria composition and alter bacterial metabolism^18–21^. These impacts may have downstream effects on glucose metabolism, such as by inhibiting the growth of *Bacteroides* species which leads to a loss of bile acid hydrolase activity and subsequent inhibition of farnesoid nuclear X (FRX) signaling^18^. Metformin-induced changes in gut microbiota are associated with increased lactate production in the intestinal lumen^22^, suggesting a role of metformin in modulating the intestinal metabolic environment. While the gut mycobiome has been implicated in T2D^3,9^, individual reports are conflicting and no previous studies have accounted for the confounding factor of antidiabetic therapies and the potential for these pharmaceuticals to directly or indirectly alter gut fungi. For example, *Saccharomyces* was relatively increased in newly diagnosed T2D compared to healthy subjects in one study ^23^ but decreased in two others ^24,25^. Increased *Candida* relative abundances in T2D was reported in two studies ^24,26^ while a third study did not report differences in *Candida*. Significant shifts in overall fungal community composition (beta diversity) compared to healthy subjects was also reported^24,25^. In these studies, most or all subjects with existing T2D diagnoses were using antidiabetic therapies, which may confound the relationship between T2D and the gut mycobiome. Furthermore, all studies had sample sizes of 14-30 people per group. Both small sample sizes and lack of stratification based on pharmaceutical use could contribute to the observed discrepancies between studies. Examining the relationship between T2D, metformin, and alterations in the gut mycobiome on a larger scale across populations is essential to better understand this potentially clinically important question.

To examine the robustness of associations between the gut mycobiome with T2D and metformin treatment, we compiled a multi-country gut mycobiome cohort from three randomized clinical trials and six cross-sectional cohorts. This resulted in a combined dataset of over 1,000 human samples with paired, individualized phenotypic data from eight studies. Our analytic methods are specifically designed to address numerous complexities of such meta-analyses including count variation, sparsity, compositional constraints, and batch effects common in these data. We then validated these results in a novel mouse study.

## RESULTS

### The most prevalent fungal genera are consistent across populations

A literature search yielded nine studies with publicly available human stool metagenomic data and individualized T2D and treatment status. The aggregated dataset comprised 1,194 samples from 963 individuals, representing three randomized clinical trials (RCTs) and three cross-sectional cohort studies that included metformin-treated individuals, in addition to three cross-sectional studies from the MetaHIT cohort containing only normoglycemic individuals (Supplementary Table S1; Figure 1A). The aggregated dataset included T2D individuals with metformin treatment (T2D-MET; n = 218), T2D individuals without metformin treatment (T2D-NOMET; n = 248), and normoglycemic individuals without metformin treatment (NORM; n = 659). Five countries were represented across Asia and Europe. A standardized bioinformatics pipeline was applied to each of the nine studies to obtain raw fastq files from sequence archives, perform quality control and host read removal, and obtain sequence count tables (see Methods). After filtering to remove very low abundant taxa, the composite dataset contained 67 fungal species comprising 43 genera that primarily represented the phylum Ascomycota, a few from Basidiomycota, and one genus from Microsporidia (Figure 1B). Analyses were conducted at the genus level to increase confidence in taxonomic assignment given the low abundance of fungal reads in shotgun metagenomics.

**Figure 1:**
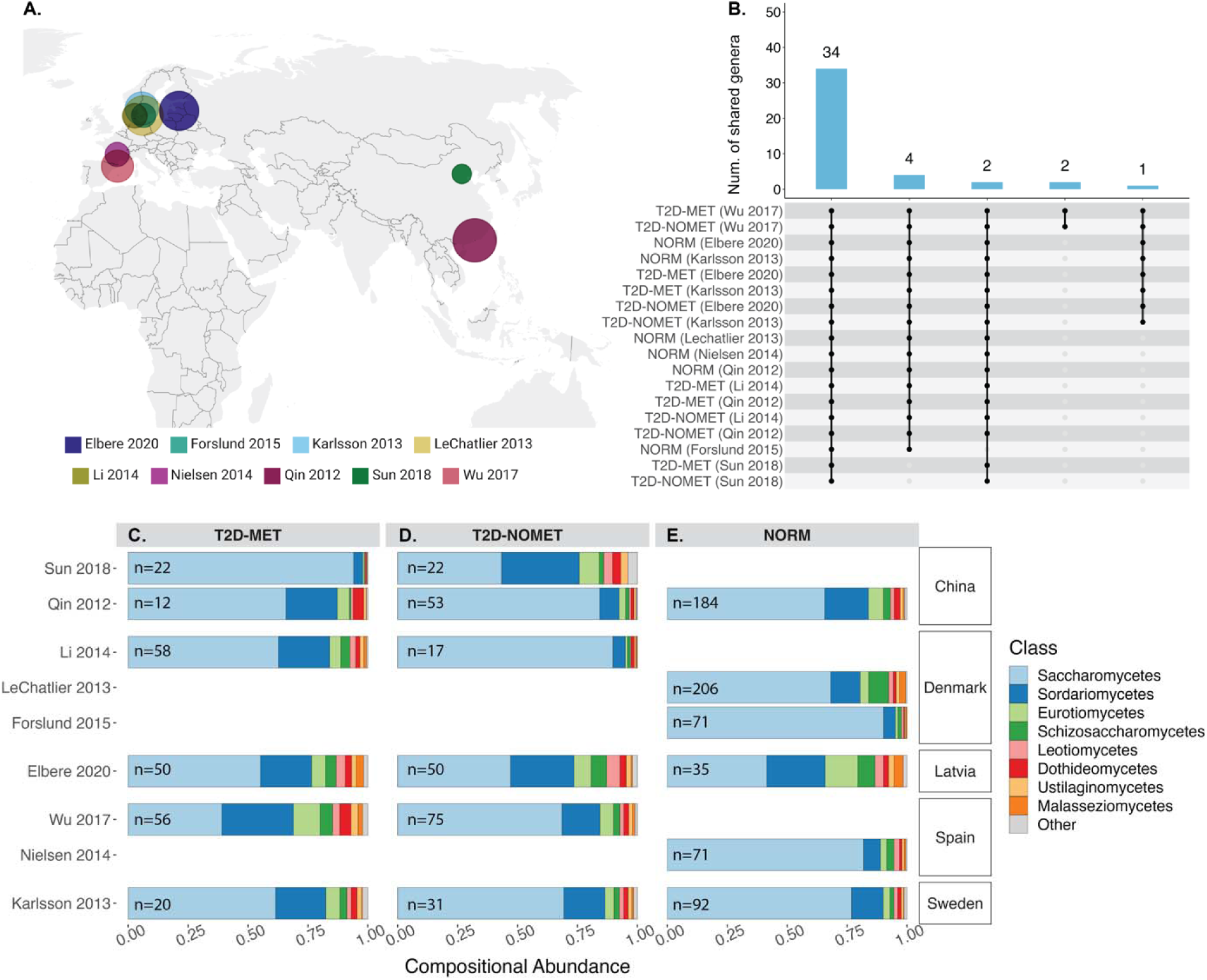
Characterization of the gut mycobiome from metagenomics data across studies. (**A**) The geographic location of each study. The size of the bubbles corresponds with total sample size. **(B)** An Upset plot demonstrating shared genera between studies. The grey dot matrix shows which study-treatment category combination shared mycobiome features (genera) and the blue bar shows the total number shared. **(C-E)** Barplots of compositional abundance at the fungal class level from each study, geographic region, and treatment category without adjustment for between-study differences.

We developed an analytical approach specifically designed to assess the reproducibility and robustness of candidate associations between gut fungi and host factors. Specifically, we used Bayesian models that have been purposely designed to address key complexities of sequence count data (e.g., sampling variation, sparsity, and compositional constraints)^30,31^ and designed our analyses to address the challenges of combining datasets (e.g., batch effects)^32,33^. We address this later challenge in two ways. First, all models fit to the combined dataset included random intercepts to minimize inter-study variation. Second, candidate associations identified from analyses of the combined dataset were further interrogated to ensure that the inferred effect was consistent when analyses were performed independently on each dataset. This later step reduces the risk that our findings are due to batch effects or outliers. Unless otherwise specified, results are stated in terms of mean Centered Log Ratio (CLR) coordinates that reflect proportional changes in fungal abundance with 95% credible intervals of the posterior distributions in brackets (e.g., mean CLR change [95% credible intervals]). Further methodological details are provided in the Methods.

Characterization of the gut mycobiome differed across studies, which was expected due to technical artifacts, like differences in sequencing depth, and geographical differences. Gut mycobiomes were largely dominated by the Saccharomycetes class which was driven by a predominance of the genus *Saccharomyces* (Figure 1C-E). However, 34 of the 43 total genera were detected in all studies (Figure 1B). Further analyses were subset to the 34 shared genera to ensure the results were not driven by presence/absence patterns that may be due to differences in sequencing depth between studies.

### Metformin treatment associates with alterations in the gut mycobiome in T2D

To determine if a relationship between metformin treatment and the human gut mycobiome emerged, we first compared the total dataset of T2D-MET (n=218) to T2D-NOMET (n=238). We detected an overall effect of metformin treatment in three fungal genera: *Fusarium* (0.147 [0.041, 0.255]; increased in T2D-MET), *Tetrapisispora* (0.230 [0.186, 0.371], increased in T2D-MET), and *Nakaseomyces* (–0.162 [–0.207, –0.036], decreased in T2D-MET) (Figure 2A). The partial R^2^ of metformin treatment (e.g., the amount of variability in the mycobiome that could be attributed to metformin) was 0.66% in the aggregate dataset, ranging from 1-10% in within-study models. Despite the variability in dosage and treatment duration, metformin had a consistently similar effect size direction and magnitude in at least four of these six studies. However, we note that while *Nakaseomyces* was found to be decreased in T2D-MET compared to T2D-NOMET in the aggregate comparison only, within-study analyses depicted higher between-study variability and conflicting directions of effect sizes for this genus, illustrated in Figure 2A.

**Figure 2:**
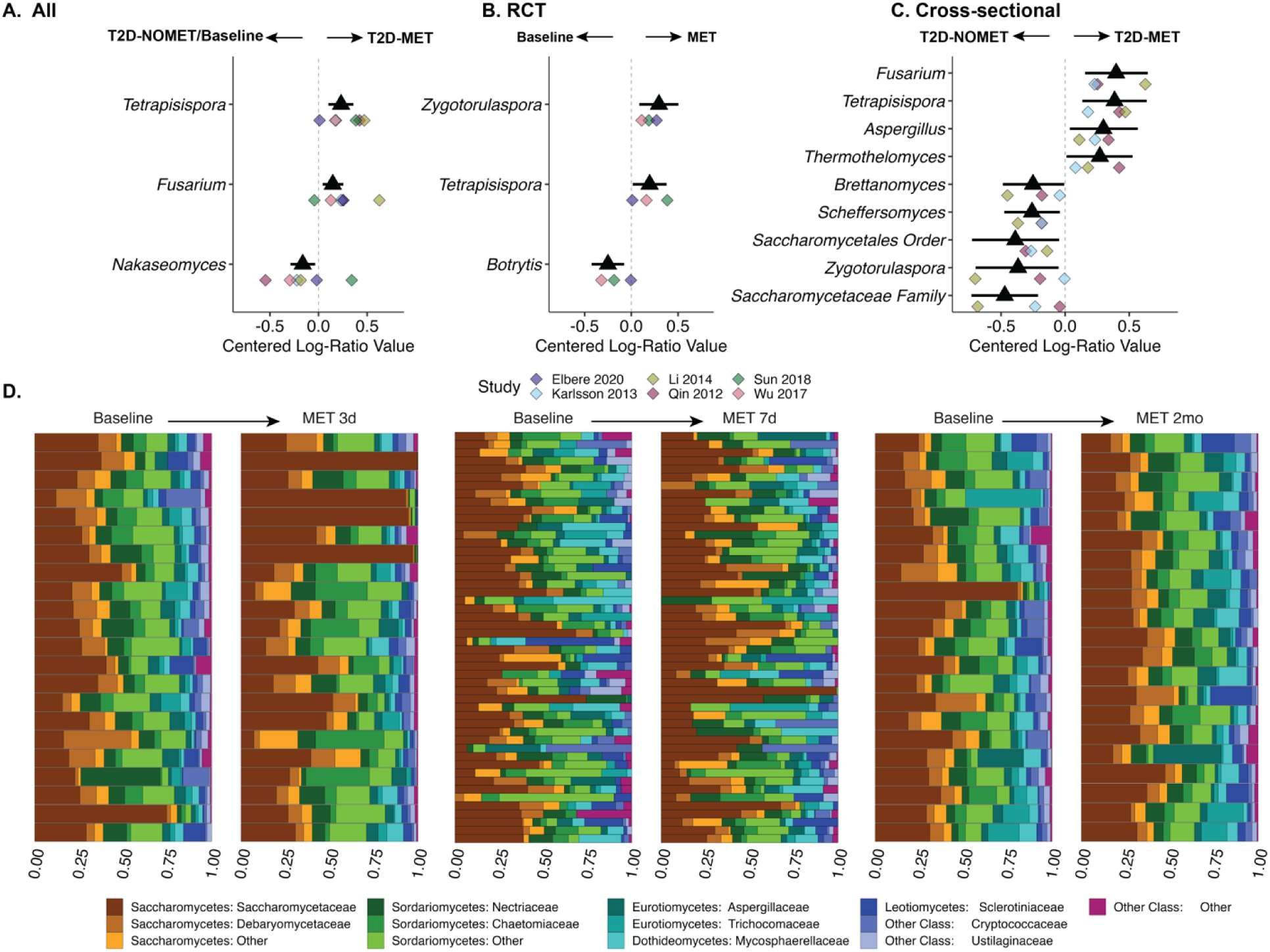
Metformin’s associations with gut fungi in T2D in (A) the total dataset, (B, D) randomized clinical trials (RCTs), and (3) cross-sectional studies. The black triangle and lines show posterior mean and 95% credible intervals of the aggregate dataset. Colored diamond points show posterior means of individual studies. The arrows show increased relative abundance at baseline or T2D-NOMET (left) or during metformin (MET) treatment (right). The dashed vertical line denotes zero association. **(D)** In three RCTs with time-series sampling, proportional relative abundance is shown at baseline and after starting metformin treatment for each study. Paired samples from the same individual are shown as neighboring horizontal bars of proportional relative abundance with the duration of metformin treatment shown above the barplot for each study. Taxa are colored as Class:Family, and less relatively abundant taxa are merged in the “other” category.

Next, we subset the dataset to further understand short-term effects of metformin treatment in newly-diagnosed, treatment-naïve T2D individuals in three RCTs (n=93). These included time-series sampling from baseline T2D diagnosis to three days, seven days, and six months following the start of metformin treatment in participants from Latvia and China. Short-term metformin treatment was associated with changes in *Zygotorulaspora* (0.298 [0.094, 0.504], increased in T2D-MET), *Tetrapisispora* (0.194 [0.013, 0.377], increased in T2D-MET), and *Botrytis* (–0.248 [–0.429, –0.074], decreased in T2D-MET) (Figure 2B). In this comparison, metformin treatment accounted for approximately 1.5% of variation (partial R^2^) in the gut mycobiome, ranging from 2-8% in within-study models. The gut mycobiomes of individual subjects had varied responses to metformin, as visualized in Figure 2D. We observed greater mycobiome variation in participants from one study that were sampled before and after receiving 2,000 mg metformin/day for three days^18^ compared to participants from two other clinical trials that received 500-2000 mg/day for 7 days^21^ or 1,700 mg/day for 2 months^19^ (Figure 2B & D). This variability in timing and dosage could contribute to study differences and inter-individual variation.

To further compare the effect of metformin treatment on fungal relative abundance in T2D, we compared three cross-sectional studies including T2D-MET (n = 90) and T2D-NOMET (n = 101) in a similar fashion. In these studies, participants could have long-term T2D and/or metformin treatment for an unknown duration and dose. Participants in this subgroup were from studies in Asia and China. Six fungal genera were different between T2D-MET and T2D-NOMET; *Aspergillus* (0.297 [0.037, 0.553]; increased in T2D-MET]), *Brettanomyces* (–0.269 [–0.488, –0.003]; decreased in T2D-MET), *Fusarium* (0.401 [0.148, 0.648]; increased in T2D-MET), *Scheffersomyces* (–0.260 [–0.478, –0.041]; decreased in T2D-MET), and two unassigned genera from the Saccharomycetales order and Saccharomycetaceae family (–0.385 [0.722, –0.056] and – 0.467 [–0.738, –0.212] respectively; both decreased in T2D-MET) (Figure 2C). In this comparison, metformin accounted for 2.8% of mycobiome variability, ranging from 3-10% within each study (partial R^2^). In these three studies, fasting blood glucose (FBG) and hemoglobin A1c (HbA1c) were similar in T2D-MET (FBG median ± inter-quartile range (IQR) 146 ± 30.6 mg/dL, HbA1c median ± IQR 6.82 ± 1.2%) compared to T2D-NOMET (FBG median ± IQR 135 ± 36.9 mg/dL, HbA1c median ± IQR 7.0 ± 3.4%) indicating that in the cross-sectional study sub-analysis, our comparison assessed the effects of pharmaceuticals and not controlled versus uncontrolled diabetes. Interestingly, some metformin-associated genera in the cross-sectional study sub-analysis differed from the genera identified in the RCT sub-analysis or full analysis. *Tetrapisispora* agreed in effect size direction and magnitude across all three comparisons.

### The mycobiome in metformin-treated T2D differs from normoglycemic individuals

To determine differences in the mycobiome of metformin-treated T2D compared to normoglycemic subjects (NORM), we compared data within three studies containing both T2D-MET (total n = 82) and NORM (total n = 262) individuals. Fungal genera that differed between T2D-MET and NORM included *Fusarium* (0.274 [0.044, 0.506], increased in T2D-MET), *Thermothielavioides* (0.222 [0.039, 0.408], increased in T2D-MET), *Cryptococcus* (0.164 [0.028, 0.307], increased in T2D-MET), and an unassigned genus from the Saccharomycetales order (–0.421 [–0.792, –0.040], decreased in T2D-MET) (Figure 3). The combination of T2D and metformin accounted for 1.3% of mycobiome variation (partial R^2^, ranging from 0.9-9% within each study). To determine the reproducibility of these findings, these candidate fungal genera were assessed in mice treated with metformin or placebo for seven days. While the metformin-treated mice were healthy and normoglycemic, we were encouraged to note similar direction and magnitude of the effect size of metformin on the mouse and human mycobiomes (Figure 3). In this controlled environment on a standard diet, metformin explained a relatively high proportion of variability in the mouse cecal mycobiome (partial R^2^ = 26.8%), suggesting that heterogeneity between experimental designs, diets, or geography may lead to an underestimation of the true treatment effect.

**Figure 3:**
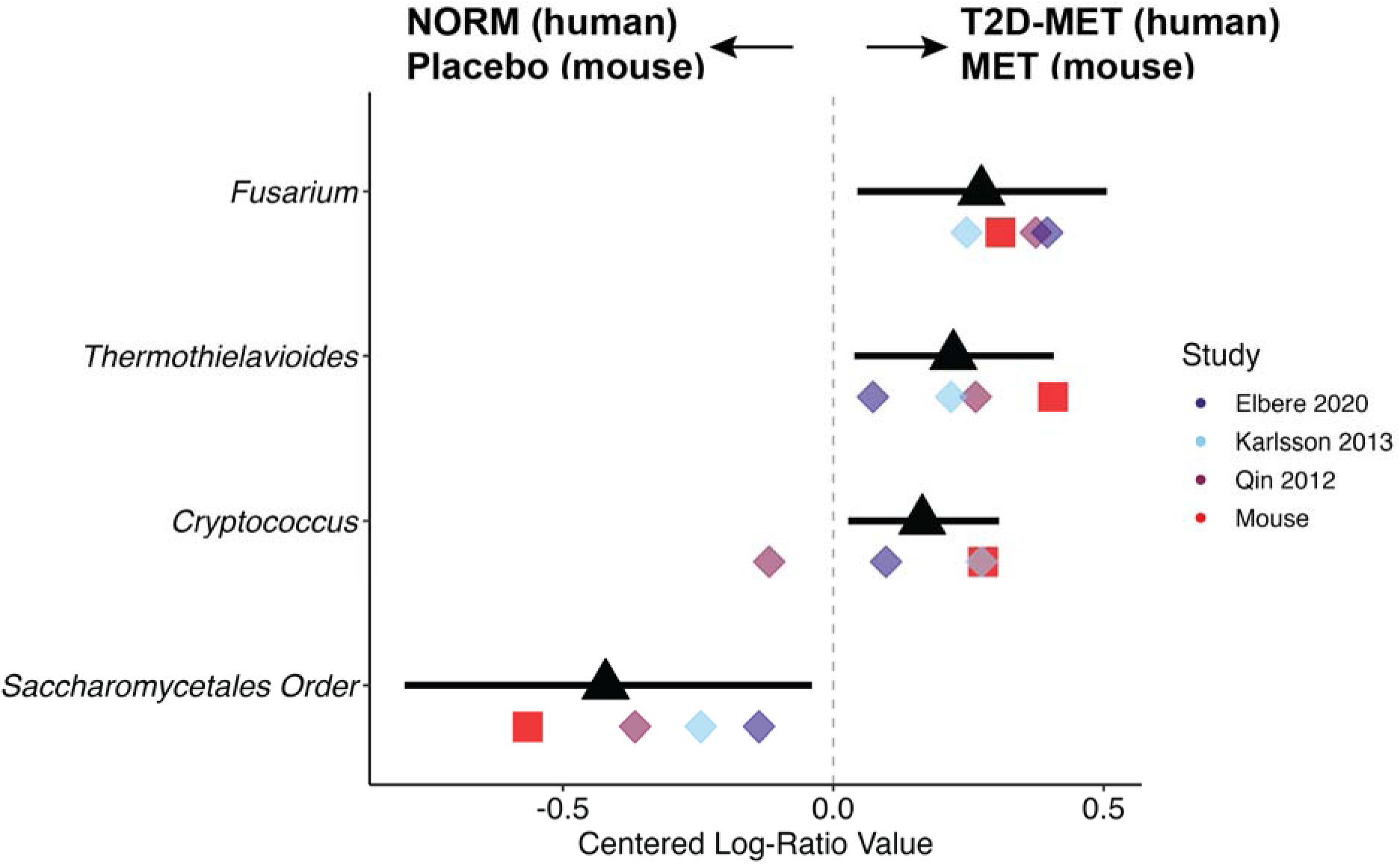
Metformin is associated with similar differences in the human and mouse mycobiomes. Black triangles and line ranges shows posterior mean and posterior 95% credible intervals of the aggregate dataset. Colored diamond points show posterior means of individual studies. Red squares show the posterior means of the mouse study. The arrows show increased abundance in NORM subjects or mice with placebo treatment (left) or T2D-MET subjects and metformin-treated mice (right). The dashed vertical line denotes zero association.

### Gut fungal relative abundance associates with clinical metrics of metabolic disease

Assessing the impact of metabolic disease on the gut mycobiome necessitated removing the confounding factor of metformin treatment. Therefore, we compared T2D without metformin treatment (T2D-NOMET; n = 134) to normoglycemic (NORM; n= 262) individuals, and additionally probed clinical indices of metabolic disease while adjusting for metformin treatment (see Methods). Shifts in fungal composition between T2D-NOMET and NORM were detected for *Saccharomyces* (0.580 [0.210, 0.959], increased in T2D-NOMET), *Nakaseomyces* (0.277 [0.085, 0.469], increased in T2D-NOMET), and *Zygosaccharomyces* (–0.434 [–0.697, –0.172], decreased in T2D-NOMET) (Figure 4A). While T2D represented approximately 1.2% of mycobiome variation in the combined dataset, within-study partial R^2^ ranged from 3-9%. To represent a wider range of metabolic phenotypes than dichotomous T2D status and increase the sample size, we leveraged individualized clinical data of fasting blood glucose (FBG; n = 784), fasting plasma insulin (FPI; n = 697), hemoglobin A1c (HbA1c; n = 815), and (body mass index, BMI; n = 766) while adjusting for metformin treatment. We found that *Saccharomyces* was positively associated and *Brettanomyces* was negatively associated with increased FBG (0.199 [0.060, 0.335] and –0.149 [–0.250, –0.045], respectively) despite variability between individual studies (Figure 4B). FPI was associated with decreased *Zygosaccharomyces* and *Tetrapisispora* (–0.098 [–0.177, –0.022] and –0.130 [–0.231, – 0.030], respectively) (Figure 4C). *Ustilago* was increased with both increased BMI and FPI (0.132 [0.786, 0.184] and 0.137 [0.083, 0.193], respectively) and *Zygotorulaspora* was associated with decreased BMI and FPI (–0.257 [–0.364, –0.148] and –0.130 [–0.231, –0.030], respectively; Figure 4C & D). None of the clinical variables were statistically associated with *Nakaseomyces* or *Fusarium* which were previously associated with metformin treatment (Figures 2 and 3). In contrast to FBG, HbA1c had very weak associations to gut fungi with high between-study variability, and no candidate genera had similar effect size direction and magnitude greater than |0.1| Centered Log-Ratio change across studies despite high within-study partial R^2^ (Table 1). The aggregated comparisons suggested that FBG, FPI, HbA1c, and BMI had low contributions to overall mycobiome variability (partial R^2^ = 0.55-0.59%), but within-study analyses demonstrated stronger relationships that ranged from 3-31%.

**Figure 4:**
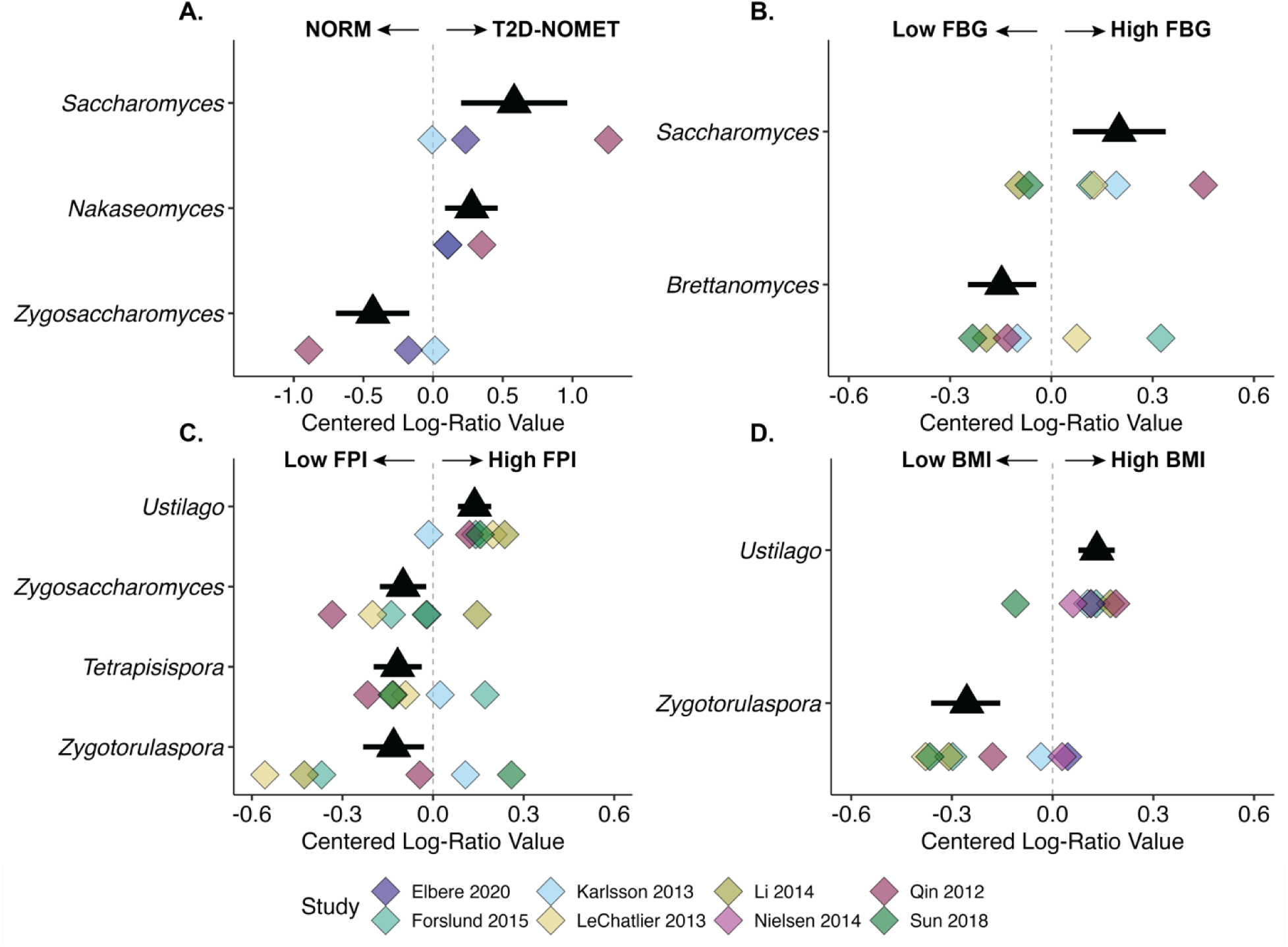
Metabolic disease is associated with a shift in genera belonging to the Saccharomycetes class. Subjects with T2D-NOMET were compared to NORM (A), and markers of metabolic disease were compared; (B) fasting blood glucose (FBG), (C) fasting plasma insulin (FPI), and (D) body mass index (BMI). In all panels, black triangles and line ranges shows posterior mean and posterior 95% credible intervals of the aggregate dataset. Colored diamond points show posterior means of individual studies. Arrows show increased fungal abundance. The dashed vertical line denotes zero association.

**Table 1:** Partial R^2^ for clinical metrics of metabolic disease calculated on the aggregate dataset and within each study. The partial R^2^ is shown with 95% credible interval in parentheses. “NA” indicates no data were available. No clinical data was available for the participants of Wu 2017. Abbreviations: FBG; fasting blood glucose, FPI; fasting plasma insulin, HbA1c; hemoglobin A1c, BMI, body mass index.

| Study | FBG | FPI | HbA1c | BMI |
| --- | --- | --- | --- | --- |
| Aggregate | 0.59 (0.34, 0.91) | 0.55 (0.33, 0.84) | 0.59 (0.37, 0.91) | 0.57 (0.34, 0.91) |
| Elbere 2020 | NA | NA | 28.4 (19.4, 38.8) | 31.5 (21.8, 42.3) |
| Forslund 2015 | 9.0 (5.9, 13.5) | 11.4 (7.4, 17.4) | 8.4 (5.2, 13.1) | 8.4 (5.4, 13.0) |
| Karlsson 2013 | 4.1 (2.4, 7.6) | 4.0 (2.5, 6.3) | 3.6 (2.1, 6.2) | 3.9 (2.2, 6.9) |
| LeChatlier 2013 | 1.6 (0.9, 2.5) | 3.0 (2.0, 4.5) | 1.8 (1.0, 3.0) | 2.2 (1.5, 3.4) |
| Li 2014 | 4.3 (2.6, 6.8) | 4.8 (3.0, 7.4) | 3.9 (2.2, 6.7) | 4.4 (2.6, 7.0) |
| Nielsen 2014 | NA | NA | NA | 5.4 (3.3, 8.4) |
| Qin 2012 | 2.8 (1.7, 4.5) | 3.1 (2.0, 4.9) | 2.6 (1.5, 4.7) | 4.5 (2.6, 7.3) |
| Sun 2018 | 11.2 (6.5, 18.2) | 11.8 (7.0, 18.9) | 10.1 (6.3, 16.4) | 11.3 (7.0, 17.8) |

## DISCUSSION

Compositional shifts in the gut mycobiome are associated with T2D, but existing studies have not accounted for the use of oral T2D pharmaceutical therapies. Given the well-established effects of metformin on gut bacterial composition and bacterial metabolism^18,19,21^, we hypothesized that metformin may confound the relationship between T2D and the gut mycobiome. We compared human subjects with T2D and metformin treatment to subjects with T2D without metformin treatment, and demonstrated effects on several fungal genera including *Fusarium*, *Tetrapisispora*, and *Zygotorulapora* that differed between RCT and cross-sectional studies. Beyond humans, the signature of metformin on *Fusarium* and *Tetrapisispora* was reproducible in a mouse study.

This analysis provides, to our knowledge, the first assessment of the effects of oral pharmaceuticals on the gut mycobiome. Antidiabetic drugs have not been examined in the context of the human gut mycobiome, but one *in vitro* experiment observed a direct interaction of metformin with the yeast *Ustilago* and postulated that metformin may extend the lifespan of microeukaryotes as it does in other eukaryotic model organisms^34^. However, the nature of this dataset precludes us from drawing any conclusions beyond associations, and the overall small contribution of metformin on gut mycobiome variability, measured as a partial R^2^, leads us to speculate that gut fungi may be indirectly affected by the metformin-induced shift in the intestinal metabolic environment ^29^.

The interactions of oral pharmaceuticals with the gut bacteriome^35^ and potentially with the gut mycobiome, as suggested in this study, necessitate accounting for these factors when assessing gut microorganisms in the context of metabolic disease. Independent of metformin treatment, T2D was associated with an increase in two genera of the Saccharomycetes class, *Saccharomyces* and *Nakaseomyces*, with a corresponding decrease of *Zygosaccharomyces*, also in the Saccharomycetes class. This was validated in the comparison of clinical markers of metabolic disease, which showed a positive correlation of *Saccharomyces* with fasting blood glucose and a negative correlation of *Zygosaccharomyces* with fasting plasma insulin. In the aggregate dataset, metformin accounted for 0.66% and T2D accounted for 1.2% of the total variability in the gut mycobiome. While this is smaller than what has been previously reported^25^, previous studies of the gut mycobiome in T2D have been conducted with marker-based sequencing of the Internal Transcribed Region, which provides deeper resolution of community diversity and is not directly comparable to shotgun metagenomics sequencing^36^. When comparing models fit to individual studies, we noticed that some studies had greater differences between T2D and NORM individuals while others had negligible differences across the board. Given the constraints of using publicly available data, we cannot exclude the possibility that these discrepancies are due to technical artifacts such as differing sequencing depths, DNA extraction techniques, sample processing, or sequencing platforms.

Our analysis represents the largest to-date gut mycobiome cohort. There is ongoing debate over what constitutes the healthy human mycobiome in addition to speculation of the true scope of intestinal fungal diversity^1,37–39^. We purposefully limited our analyses to the most abundant and prevalent fungal genera to exclude minor taxa that may be determined by environmental exposures or may even be spurious taxonomic assignments. For example, the two least prevalent fungi in our aggregated dataset, two unassigned genera from the families Chaetomiaceae and Mycosphaerellaceae, were detected in only a few individuals from one study and have not been previously associated with the human gut mycobiome. The most abundant fungi across the dataset were prevalent in approximately 95% of subjects, suggesting a ubiquitous presence regardless of diet or geography, and were similar to previous reports that confirmed the presence of 34 fungal genera with culture-dependent approaches^39^.

This analysis is limited by the constraints of using publicly available data, as laboratory controls such as DNA extraction blanks and clinical metadata such as dietary habits and additional pharmaceutical consumption are simply not available. Dietary recommendations are standard of care for newly diagnosed T2D, and thus it is possible that some differences we attributed to T2D could be due to dietary changes^40^. As fungi comprise a low proportion of total gut microbial biomass^4^, the detection of fungi in shotgun metagenomics reads can be difficult. We took a simplistic approach with kmer-based taxonomic assignment using similar methods to existing mycobiome surveys^28,41,42^. The pitfalls of this bioinformatics approach include a risk of false positives that inflate the dataset or detection of contaminants as true taxa. We minimized these risks by first removing human host reads and reads that aligned to a bacterial & archaeal database, and then comparing the unaligned reads to the fungal database, and further filtered out very low abundant taxa with the *a priori* assumption that they may be spurious assignments or contaminants. The Bayesian Multinomial Logistic-Normal modeling approach ensured that between-study variation was minimized, and our priors were conservatively chosen to assume there were no true differences between comparison groups. In addition to that approach, we further confirmed candidate taxa identified in the aggregated dataset were not driven by outliers by ensuring those taxa were consistently different across multiple independent studies and with similar direction. Given the large amount of variability in populations, experimental design, and sample preparation methods between the included studies, the consistent reproducibility of our results and replication in a controlled murine study suggests non-trivial associations of some gut fungi with T2D and metformin treatment.

In summary, we identify, for the first time, associations between the gut mycobiome and oral pharmaceutical treatment. We show distinct fungi associated with T2D and with metformin and validated the changes in those fungi in a murine study with metformin administration. Metformin and T2D generally accounted for less than 5% of total mycobiome variation in the aggregate comparisons, suggesting a subtle but significant effect of both metformin and T2D. This analysis provides a framework by which to evaluate shifts in gut microbes across large datasets using analyses specifically designed for the challenges of compositional sequencing data. These analyses across larger sample sizes and heterogeneous populations are paramount to provide validation of findings reported in smaller, isolated human studies. Our findings highlight the need to consider the potentially confounding effects of oral pharmaceuticals and other exposures when considering linkages of gut microbes and human diseases. Future research should seek to integrate data across gut bacteria, fungi, and other microbial inhabitants of the human gastrointestinal tract to further understand the contributions of all microbial life to human gut homeostasis.

## METHODS

### Metagenomics acquisition and bioinformatics

To obtain metagenomics data, a literature search was performed, and relevant articles were screened for the following criteria: human studies, collection of stool samples for shotgun metagenomics sequencing, and type 2 diabetes or metformin treatment. Three additional datasets from the MetaHIT cohort were included to increase the sample size of normoglycemic comparisons (accessions PRJEB5224, PRJEB4336, PRJEB1220). Duplicate participant IDs between the three accession numbers were removed, and the MetaHIT cohorts retained the label of the study they first appeared in. Thus, nine studies were included in the analysis comprising 1,194 samples (Supplementary Table S1).

Metagenomic data was retrieved in fastq format and re-analyzed under one standard pipeline to minimize between-study variability. None of the studies provided sequences from technical controls. Metagenomic data was first downloaded from NCBI Sequence Read Archive or European Nucleotide Archive with Entrez Direct software ^43^. Adapter contamination was removed with fastp v0.20.1 and human reads were removed with bowtie2 v2.4.2 aligning to the human genome version HG37v0.1 ^44,45^. Quality and length trimming was performed in fastp to discard reads with a quality score in a 4-base sliding window under 20 and length under 2/3 of the average read length. Bacterial taxonomy was assigned by mapping to the standard kraken2 v2.1.1 database build ^46^. The unassigned reads were then mapped to the standard fungal library build in kraken2, which comprises the complete RefSeq fungal genomes and proteins. Both databases were retrieved in November 2021. Kraken2 reports were converted to biom tables with kraken-biom (https://github.com/smdabdoub/kraken-biom). Further analyses were conducted in R software v4.0.13 ^47^. A filtering approach was undertaken to remove very low abundant taxa. Filtering was applied within each study to account for differences in sequencing depth. Taxa that were unassigned at the phylum level and taxa detected in fewer than 10% of samples within a study were removed, and taxa were further filtered to remove species with fewer than 0.001% of total assigned reads. Samples with zero fungal reads were removed (n = 7, all from Qin et al 2012).

Clinical and phenotypic metadata were collected from NCBI’s Sequence Read Archive metadata tables, the study’s supplementary materials, or data provided by the study authors through institutional Data Transfer Agreements. Metadata was only collected if individualized data was provided and if the data could be matched back to the archived sequencing sample through a provided sample name. Metformin and T2D status were assigned to each sample based on the categorization given in the study metadata.

### Bayesian multinomial logistic-normal linear regression modeling

To model the effect of these covariates on fungal and bacterial composition, we used a Bayesian multinomial logistic-normal linear regression model ^48^. We selected this model since it accounts for sampling variation in the observed counts and satisfies compositional constraints. This model is efficiently implemented via the pibble function in the R package *fido* ^48^. This structure models the log ratio coordinates using a Bayesian linear regression model. The structure for each model was dependent on the scientific question; covariates used for each model in the main text are defined in Table 2. In global comparisons that included time-series data from randomized clinical trials, a random intercept was added for each individual. To avoid misattributing variability to study differences, global models contained only matched data from each study; that is, instead of comparing all T2D-MET individuals to all NORM individuals, we only used data from studies that included both T2D-MET and NORM.

**Table 2:** Bayesian MLN model covariates. All models except the mouse data (marked with *) were fit to the aggregate dataset and further validated with individual study models. Individual study models were identical to the global model without the *[x _j_,(study)]* covariate. *[x_k_…,(individual)]* represents a random intercept for each individual in time-series designs.

| **Comparison** | **Figure** | **Covariates** |
| --- | --- | --- |
| T2DMET - T2DNOMET; All | Figure 2A | $[1, x_{j,(study)}, x_{j,(metformin)}, x_{k,...,(individual)}]$ |
| T2DMET - T2DNOMET; RCT | Figure 2B | $[1, x_{j,(study)}, x_{j,(metformin)}, x_{k,...,(individual)}]$ |
| T2DMET - T2DNOMET; Cross-sectional | Figure 2C | $[1, x_{j,(study)}, x_{j,(metformin)}]$ |
| T2DMET - NORM; Humans | Figure 3 | $[1, x_{j,(study)}, x_{j,(metformin)}]$ |
| T2DMET - NORM; Mouse* | Figure 3 | $[1, x_{j,(metformin)}, x_{k,...,(individual)}]$ |
| T2DNOMET - NORM | Figure 4A | $[1, x_{j,(study)}, x_{j,(T2D)}]$ |
| Fasting blood glucose (FBG) | Figure 4B | $[1, x_{j,(study)}, x_{j,(metformin)}, x_{j,(FBG)}]$ |
| Fasting plasma insulin (FPI) | Figure 4C | $[1, x_{j,(study)}, x_{j,(metformin)}, x_{j,(FPI)}]$ |
| Hemoglobin A1c (HBA1C) | - | $[1, x_{j,(study)}, x_{j,(metformin)}, x_{j,(HBA1C)}]$ |
| Body mass index (BMI) | Figure 4D | $[1, x_{j,(study)}, x_{j,(metformin)}, x_{j,(BMI)}]$ |

Since we chose a Bayesian regression model, we set four specific hyperparameters — Γ, Θ, Ξ, and υ — for model fitting. We set Γ to be a Q dimensional identity matrix where Q is the number of covariates in a given model. This reflected a weak prior assumption that the covariates are uncorrelated. Additionally, we set Θ to be a (D − 1) ×Q matrix of zeros where Q is defined as before and D denotes the number of taxa. This reflects a prior belief of a null relationship in the additive log-ratio space (ALR) between each covariate and taxa. Furthermore, we chose our values of Ξ and υ to reflect a weak prior belief that each taxon is uncorrelated in terms of absolute abundances. Thus, we chose υ = D + 3 and selected Ξ such that it corresponded to a D-dimensional identity matrix in the space of log abundances scaled by a factor of (υ −D). This corresponds to a (D− 1) × (D− 1) matrix with ξii = (υ −D) and ξi j = (υ −D)/2. These specifications were checked using prior predictive checks available in *fido*. Missing values in each model were removed. We simulated 5,000 posterior draws for each model. The function in *pibble* natively fits these models in terms of ALR coordinates. After model fitting, we transformed the resultant parameters to centered log ratio (CLR) coordinates using the *fido* function to_clr. For each parameter of interest, we calculated posterior means and 95% credible intervals. The resulting intervals were assessed for whether they contain zero; parameters for which these intervals did not cover zero were investigated further. This model was fit to the aggregated dataset for each comparison, and candidate genera identified as having 95% credible intervals outside of zero were subjected to further filtering. These candidate genera were assessed in models fit to each study and only retained if the majority of studies agreed in the effect size direction (positive or negative fold change) and putatively non-trivial magnitude (Centered Log-Ratio change > |0.1|). This filtering approach ensured that the aggregate comparisons were robust to between-study differences or outliers. Due to the model structure, partial pooling of the estimates occurs and multiple hypothesis corrections are unnecessary ^31,49^.

### Animal Study

Eight-week-old C57Bl/6N mice on a standard diet were treated with normal water (placebo, n=6) or 2 mg/mL metformin in drinking water (n=6) for seven days. Mice were maintained in a specific pathogen-free (SPF) environment, and all animal protocols for mouse experimentation were reviewed and approved by the National Cancer Institute (NCI) Animal Care and Use Committees of National Institutes of Health (NIH). On day 7, the mice were fasted for 3 hours prior to euthanasia and cecal contents were collected. Genomic DNA was extracted from cecal contents with MagMax reagents (ThermoFisher Scientific) in an automated Kingfisher instrument following manufacturer instructions. One extraction blank and one mock community (ZymoBiomics Microbial Community Standard) were included in DNA extraction and carried through sequencing. Extracted DNA quantity was quantified on a Qubit and genomic DNA was shipped overnight on dry ice to Novogene Corporation (Sacramento, California, USA) for shotgun metagenomics sequencing on the Illumina NovaSeq platform, resulting in 150×150 bp paired-end reads. The shotgun metagenomics data was subjected to an identical bioinformatics pipeline as described above except reads were aligned against the mouse genome build GRCm39 (GCF_000001635.27) for host read removal. Taxa not assigned at the phylum level were removed, and putative contaminants based on the blank control were detected with the decontam R package. A brief report of the control samples is presented in Supplementary Figure 1.

### Data Availability

The metagenomic data generated from the mouse study is available at the NCBI Sequence Read Archive under accession PRJNA940065. The accession codes of archived data obtained in this study are PRJNA486795, PRJEB1786, PRJEB39500, PRJEB2054, PRJEB4336, PRJEB5224, PRJEB1220, PRJNA422434, and PRJNA361402. Individualized phenotypic data is available in each study’s supplementary material or the corresponding SRA repository. Individualized clinical data from two studies (Elbere 2020 and Sun 2018) were obtained from study authors or through an institutional Data Transfer Agreement. Source code that generated all analyses and figures are available at https://github.com/gandalab/metformin-mycobiome.

## Acknowledgements

We would like to thank Dr. Istvan Albert for his guidance in modifying bash scripts originally provided in the Biostar Handbook. Computations for this research were performed on the Pennsylvania State University’s Institute for Computational and Data Sciences’ Roar supercomputer. We are also grateful to the participants of these studies and the study investigators who made the data available.

