## Supplementary Materials for "Changes in the Type 2 diabetes gut mycobiome associate with metformin treatment across populations"

**Supplementary Table S1: Summary of included studies.** The study first author and year is shown with the Sequence Read Archive accession number, experimental design, and sample size. Sample sizes are represented as total number of individual human participants and total number of samples. The RCTs had time-series designs, resulting in multiple samples per participant and greater sample n than individual n.

| *Author* | *Accession* | *Participant n* | *Sample n* | *Experimental Design* |
|---|---|---|---|---|
| Elbere 2020 | PRJEB39500 | 85 | 203 | RCT: T2D-MET, T2D-NOMET, NORM, NORM-MET |
| Forslund 2015 | PRJEB2054 | 71 | 71 | CS: NORM |
| Karlsson 2013 | ERP00246 | 143 | 143 | CS: T2D-MET, T2D-NOMET, NORM |
| LeChatlier 2013 | PRJEB4336 | 207 | 207 | CS: NORM |
| Li 2014 | ERP004605, PRJEB5224 | 75 | 75 | CS: T2D-MET, T2D-NOMET |
| Nielsen 2014 | PRJEB1220 | 71 | 71 | CS: NORM |
| Qin 2012 | PRJNA422434 | 249 | 249 | CS: T2D-MET, T2D-NOMET, NORM |
| Sun 2018 | PRJNA486795 | 22 | 44 | RCT: T2D-MET, T2D-NOMET |
| Wu 2017 | PRJNA361402 | 40 | 131 | RCT: T2D-MET, T2D-NOMET |
| | ***Total*** | ***963*** | ***1,194*** | |

Abbreviations: RCT, randomized clinical trial; CS, cross-sectional cohort study.

**Supplementary Figure 1: Mock community and blank controls were extracted for DNA alongside cecal contents and carried through sequencing in the murine study.** After pre-processing through an identical bioinformatics pipeline as the human dataset, the mock community and blank controls are shown. Fungal and bacterial composition of the mock and blank controls is shown in (A, C) fractional relative abundance, and (B, D) absolute counts, respectively. Both yeasts expected in the mock community, *Saccharomyces* and *Cryptococcus*, were detected and comprised almost all relative abundance, while the blank sample contained fewer than 50 reads of contaminant *Fusarium*.

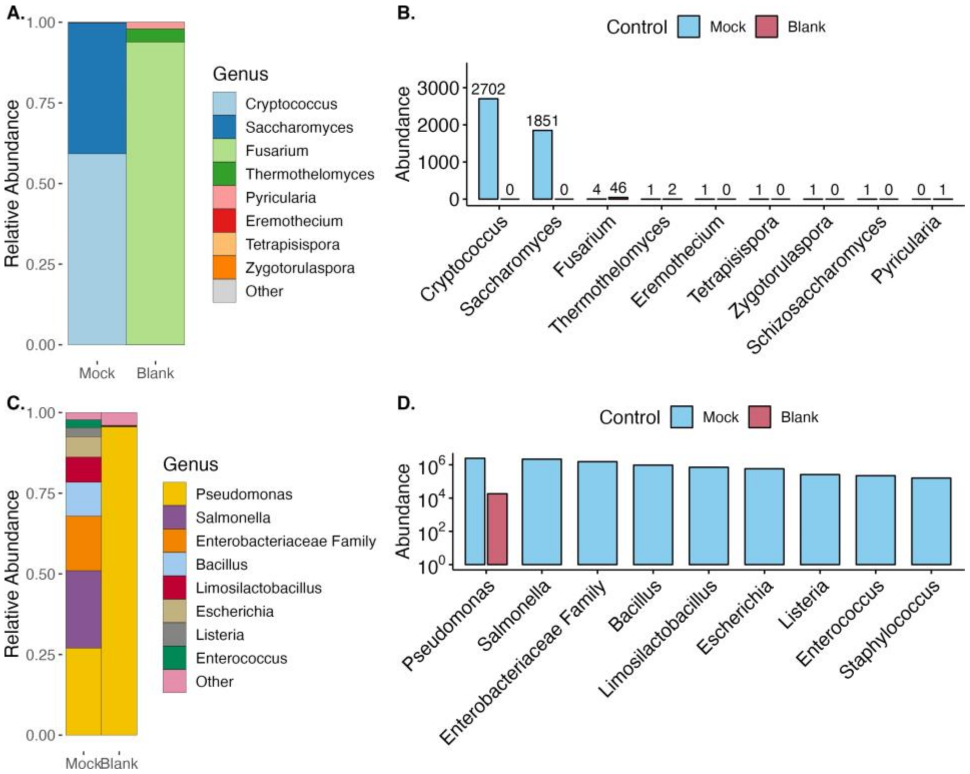
